# Establishing an auxin-inducible GFP nanobody-based acute protein knockdown system to mimic hypomorphic mutations during early medaka embryogenesis

**DOI:** 10.1101/2025.05.23.655727

**Authors:** Kaisa Pakari, Sevinç Jakab, Encarnación Sánchez Salvador, Christian Thiel, Joachim Wittbrodt, Thomas Thumberger

**Author notes:** Weill Institute for Neurosciences, Memory and Aging Center, University of California, San Francisco, 94158 San Francisco, USA.

## Abstract

Creating hypomorphic mutations are crucial to study gene function *in vivo*, especially when null mutations result in (embryonic) lethality. This is especially the case for enzymes involved in glycosylation that, when mutated in human patients, are causing the disease congenital disorders of glycosylation (CDG). To resemble the patient conditions, it would be ideal to acutely modulate the proteins in question to directly interfere with protein levels of such essential enzymes. These methods offer to establish pathogenic enzyme levels resembling net enzyme activity reported from patients suffering from CDG, with Phosphomannomutase 2 - CDG (PMM2-CDG) as the most common form.

We established an auxin-inducible acute protein knockdown system for the use in the teleost fish medaka (*Oryzias latipes*) by combining an improved degron (AID2) technology with a mAID-nanobody targeting endogenously GFP-tagged Pmm2 protein. We generated a fishline expressing a functional Pmm2-GFP fusion protein, by single copy integration of *GFP* into the *pmm2* locus. Upon induction, the degron system efficiently reduced Pmm2-GFP levels and enzyme activity, recapitulating the activity level of the hypomorphic mutations associated with PMM2-CDG in patients. This broadly applicable approach enables the investigation of CDG disease mechanisms during early embryonic development through reduction of protein abundance mimicking hypomorphic mutations and thus substantially expands the range of the genetic toolbox.

**Summary Statement:** The combination of TIR1F74G and mAID-GFP-nanobody enables efficient acute knockdown of endogenously GFP-tagged proteins in medaka. This approach successfully reduced Pmm2 enzyme activity to pathological levels as seen in PMM2-CDG patients.

## Introduction

Multisystemic diseases are often caused by mal-functioning proteins, which result from genetic mutations that impact on enzyme activity. Animal models are crucial to understand the molecular mechanisms underlying disease initiation and progression, and offer to evaluate potential therapeutic treatments of such diseases. However, generating viable animal models by precisely recreating such mutations is facing multiple challenges: targeting essential genes with the available genome editing toolkits causes unwanted insertion and deletions or coding sequence changes that are often lethal for the genome edited organism. This makes creating, identifying and breeding hypomorphic alleles nearly impossible. Further, stable genetic lines can only be maintained in a heterozygous state, i.e. due to Mendelian genetics, only a quarter of the offspring will be homozygous and may develop the (rare disease) phenotype.

An alternative and more direct approach is the manipulation of protein abundance which allows to mimic hypomorphic conditions in a controlled manner. This is especially of importance for studying rare human diseases such as congenital disorders of glycosylation (CDG). This rare group of complex metabolic disorders is caused by disrupted glycosylation, a critical post-translational modification process. CDG patients usually harbor hypomorphic mutations which reduce the enzyme activity and compared to non-viable null mutations cause less severe but often systemic symptoms. The most prevalent form of CDG is caused by mutations in the *Phosphomannomutase 2* (*PMM2*) gene (Ng et al., 2024) encoding the enzyme that converts Mannose-6-phosphate to Mannose-1-phosphate, the initial step for all ER-based glycosylation (*N*-glycosylation, *C*- and *O*-mannosylation, GPI anchor biosynthesis) (Sharma et al., 2014). Most of the reported PMM2-CDG patients have a clearly reduced PMM2 enzyme activity (Pajusalu et al., 2024). Various cell line models and studies including patient derived fibroblasts that carry hypomorphic *PMM2* mutations have helped to acquire the basic understanding of PMM2 activity on the cellular level. Available organismal models that are based on genetic PMM2 mutations either show a subset of patient phenotypes or suffer from early lethality (Chan et al., 2016; Cline et al., 2012; Himmelreich et al., 2015; Parkinson et al., 2016a, 2016b; Schneider et al., 2012; Thiel et al., 2006; Zhong & Lai, 2025).

In contrast, controlled reduction of PMM2 protein abundance would affect all treated specimens and will result in reduced net enzyme activity comparable to the hypomorphic conditions seen in patients. Progressive diseases like PMM2-CDG are characterized by an early onset of phenotypes which is challenging to study in mammals due to the intrauterine development of the embryo. We thus turned to the teleost fish medaka (*Oryzias latipes*), a well-suited model for studying human diseases and related gene functions (Doering et al., 2023; Gücüm et al., 2021; Hammouda et al., 2021), as the early embryogenesis can be thoroughly examined due to the extrauterine development and transparency of the embryos (Wittbrodt et al., 2002).

As an alternative to recreating patient-specific alleles, controlled reduction of the PMM2 protein abundance can recreate patient conditions. An efficient method for acute and inducible reduction of PMM2 activity at the organismal level, targeted protein degradation using degron systems would provide the most direct approach.

Degron systems allow fast degradation of proteins-of-interest via the ubiquitin-proteasome pathway in a highly specific, inducible and reversible manner. The targeted inactivation of proteins can be achieved by the auxin-inducible degron (AID) system, an E3 ubiquitin ligase complex, in which the plant F-box protein TIR1 (transport inhibitor response 1) dimerizes with AID-tagged proteins in the presence of the plant hormone auxin (Nishimura et al., 2009). This dimerization recruits the tagged proteins to the endogenous Skp1–Cul1–F-box (SCF) E3 ligase complex and results in ubiquitination and degradation of the tagged protein. A drawback of the canonical TIR1 protein is its basal activity in the absence of auxin (Natsume et al., 2016). Improved degron systems differ between TIR1 F-box variants employed (Nishimura et al., 2020; Yesbolatova et al., 2020). Another limiting factor for these approaches is the requirement of tagging the protein-of-interest with AID. To gain flexibility and foster targeted line generation with a visible selection readout, we used a deGradFP approach in which GFP-tagged proteins are recognized by a highly GFP-specific nanobody (vhhGFP4) (Caussinus et al., 2011; Caussinus & Affolter, 2016) fused to a minimized mAID (Daniel et al., 2018). Successful degradation is thus visible due to the loss of fluorescence linked to target protein depletion upon induction.

Thus, generation of a stable Pmm2-GFP fusion line in medaka not only allows for acute interference with a visual readout, but additionally provides a systemic expression analysis of Pmm2 *in vivo*.

Here we combined a specificity-improved auxin-inducible degron system (TIR1(F74G)) with a mAID-nanobody targeting GFP-tagged protein approach for the use in medaka. We generated and functionally validated an endogenously tagged Pmm2-GFP medaka line. Furthermore, we used a visible readout to demonstrate the efficacy of the combinatorial degron system by interfering with Pmm2-GFP levels and biochemically confirmed the reduced net enzyme activity of Pmm2 following induction of the degron system. We established a flexible system to manipulate Pmm2 levels and activity in a specific and efficient manner, providing a valuable tool for further studies on the role of Pmm2 in development.

## Results and Discussion

For the application of degron systems in GFP-tagged lines, it is essential that the fluorophore does not interfer with the funcionality of the fused protein and that both poly-peptides remain stably bound. We used CRISPR/Cas9 to trigger the homology directed repair (HDR) mechanism inserting the coding sequence of a flexible linker and *GFP* at the C-terminus of the endogenous *pmm2* locus, yielding a seamless Pmm2-GFP fusion (Fig. 1A). To increase single integration efficiency and enhance HDR, the PCR-donor sequence was protected with biotin at the 5’ end of each primer during amplification (Gutierrez-Triana et al., 2018). Following injections at the 1-cell stage, GFP fluorescence was used as a proxy for successful protein targeting. Offspring descending from positively screened embryos were validated by genotyping and raised. In the validated stable line we established, Pmm2-GFP expression was following the endogenous expression and was found ubiquitously and continuously (Fig. 1B, Fig. S1). Homozygous animals exhibited normal growth, development, and fertility, indicating that the GFP-tag did not interfere with Pmm2 function. A PMM2-enzyme assay performed on wild-type and Pmm2-GFP homozygous offspring revealed comparable enzyme activity, with a mean specific activity of 0.093 U/mg for wild-type and 0.095 U/mg for Pmm2-GFP (n total = 93 of three independent replicates, respectively; Fig. 1C; (Korner et al., 1998)). Single copy integration of *GFP* into the *pmm2* locus was confirmed by PCR-genotyping and Southern blot analysis (Fig. S1). Sequencing of the *pmm2* cDNA reverse transcribed from mRNA isolated from the stable line confirmed the correct sequence and splicing of the transcript, resulting in a single open reading frame and seamless fusion of *pmm2* and *GFP* coding sequences in mature mRNA. This properly fused mRNA was transcribed and spliced from a modified genomic locus with a partial duplication of the left homology flank (LH) (i.e., the entire coding sequence of exon 8 (ENSORLE00000306983) and parts of the leading intron). The right homology flank (RH) of the donor was seamlessly integrated via HDR (Fig. 1A, Fig. S1). Western blot analysis further confirmed the Pmm2-GFP fusion protein as the sole product from the tagged locus, without additional individual Pmm2 or GFP proteins (Fig. 1D).

**Figure 1:**
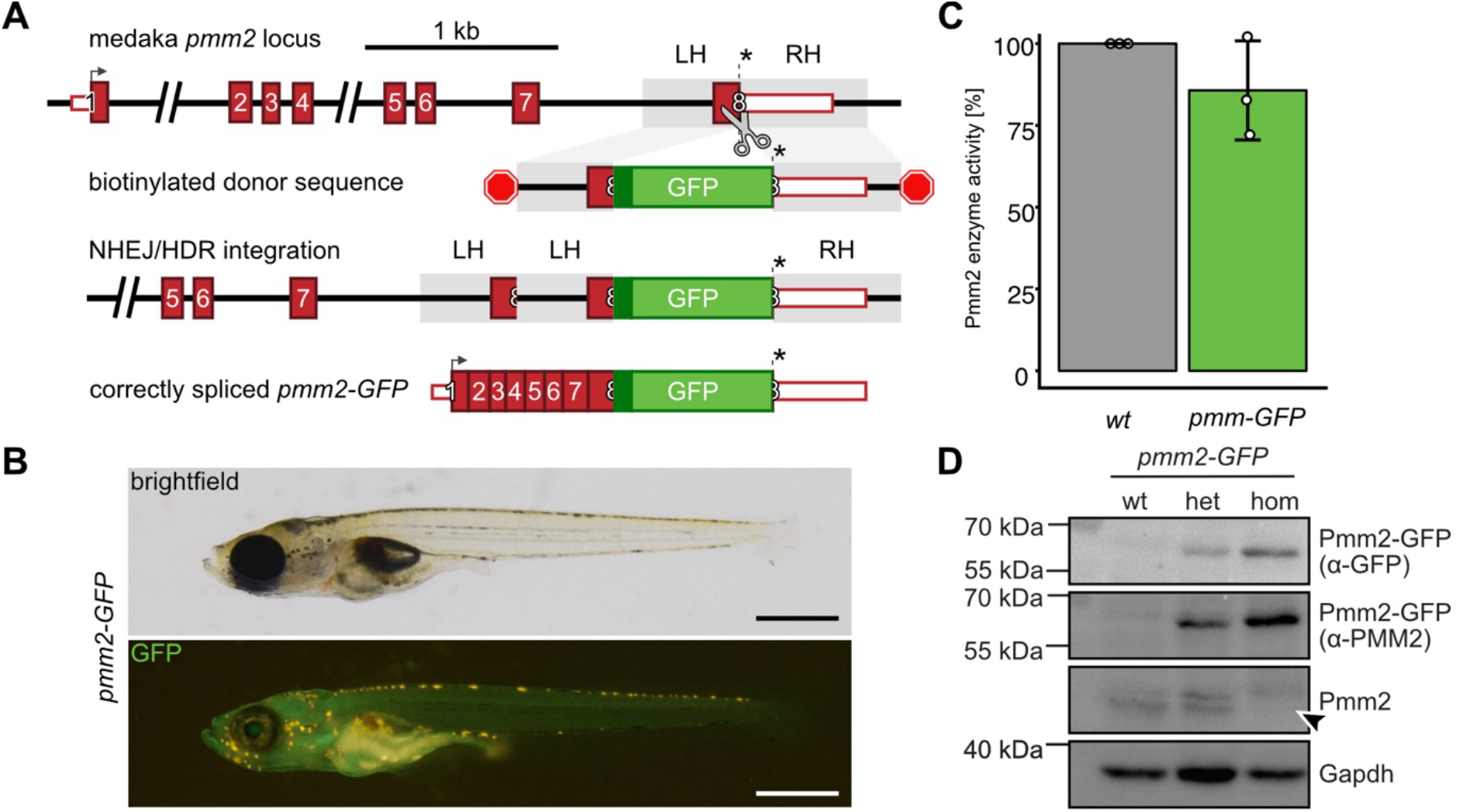
Endogenous *GFP*-tagging of *pmm2* does not interfere with Pmm2 function in medaka. A) Schematic representation of the medaka *pmm2* gene locus. Endogenous GFP knock-in outline at the C-terminus of the *pmm2* coding sequence via CRISPR/Cas9 (scissors) and biotinylated (red stop signs) donor template (coding sequence, red boxes; untranslated region (UTR), white boxes. Southern blot analysis (Fig. S1) revealed a single copy integration by homology directed repair (HDR) with the right homology flank (RH) and a non-homologous end-joining (NHEJ) event of the left homology flank (LH), leaving a duplication of the splice acceptor and coding sequence of exon 8. Transcript analysis revealed correctly spliced *pmm2-GFP* mRNA (Fig. S1). B) Representative hatchling homozygous for *pmm2-GFP* shows ubiquitous GFP expression and proper development. C) Biochemical enzyme activity assay highlights matching levels of wild-type (wt) and GFP-tagged Pmm2 (pools of 25 embryo lysates). D) Western blot analysis of lysates of wild-type (wt), Pmm2-GFP heterozygous (het) and Pmm2-GFP homozygous (hom) hatchlings, reveal the presence of Pmm2-GFP as a stable fusion protein using Pmm2 and GFP specific antibodies; see absence of Pmm2 band in hom hatchlings (arrowhead), Gapdh was used as loading control. Scale bar 50 µm. *, stop-codon; cDNA, complementary DNA; gDNA, genomic DNA; het, heterozygous; ol, *Oryzias latipes*; UTR, untranslated region

### The TIR1/mAID-GFP-nanobody degron system is effective but harbors high basal activity in medaka

The combination of F-box protein OsTIR1 with nanobodies for degradation of GFP-fusion proteins in an auxin inducible manner was previously developed in human cell culture lines and applied in zebrafish embryos (Daniel et al., 2018). Here we attempted to target the endogenously tagged Pmm2-GFP with the vhhGFP4 nanobody to recruit it to Tir1 in an auxin inducible manner, thereby facilitating ubiquitination by the SCF complex (Fig. 2A). As the tolerated concentration of chemical effectors differs between organisms, an auxin (1-Naphthaleneacetic acid; NAA) toxicity test was first performed on wild-type medaka embryos. Embryos were subjected to a range of different NAA - concentrations and phenotypes were assessed (Fig. S2). Embryos at high NAA concentrations (≥ 5 mM) displayed a wide range of developmental defects including blood clotting, edema around the heart or overall embryonic misdevelopment (Fig. S2). NAA incubations up to 0.5 mM did not lead to any scorable phenotypes.

**Figure 2.**
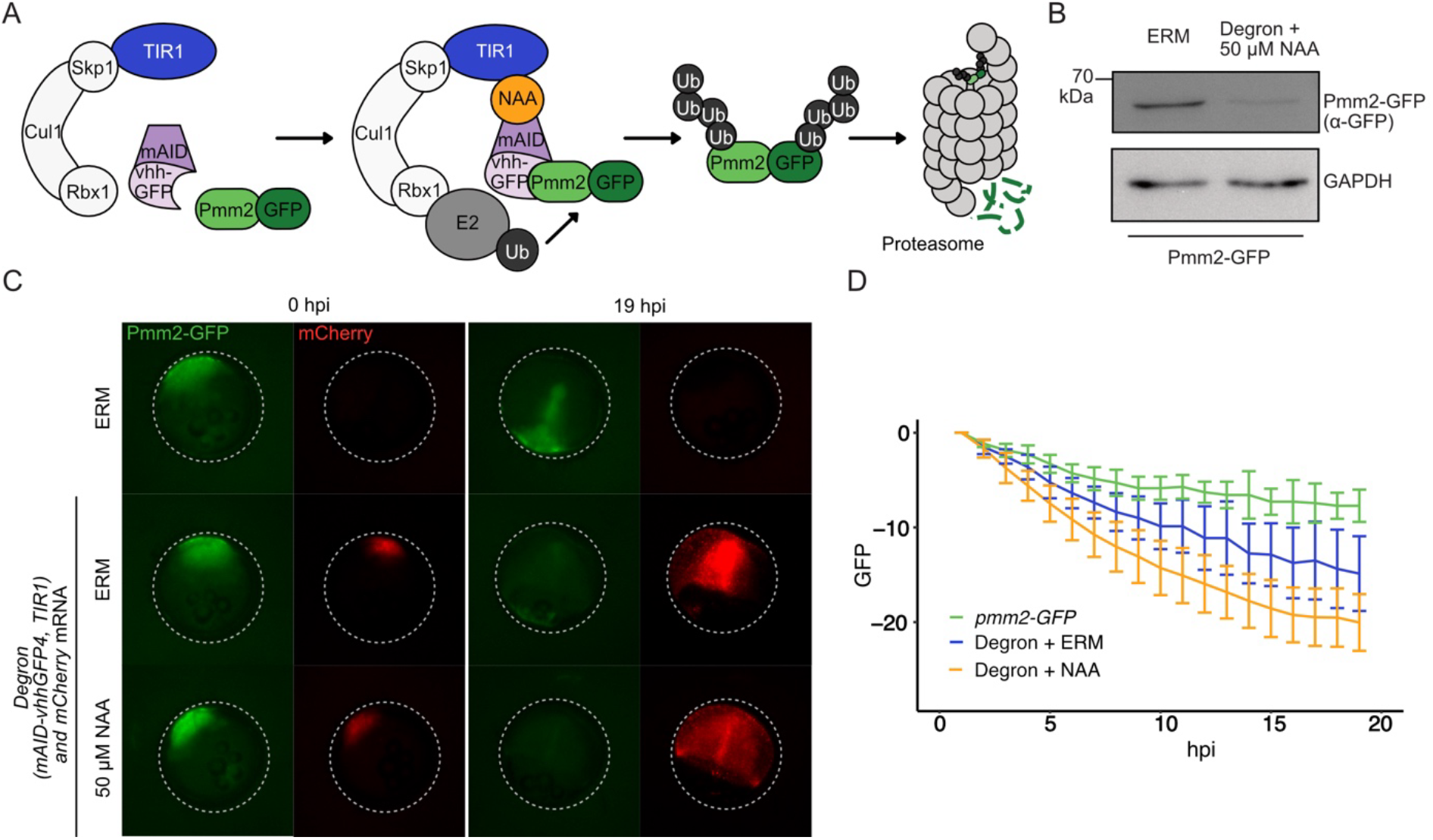
TIR1 and mAID/GFP-nanobody degron system has strong basal activity in the absence of auxin in medaka. A) Schematic representation of a mAID and GFP-nanobody based, auxin inducible degron system (Daniel et al., 2018) planned for acute degradation of Pmm2-GFP in medaka. Upon auxin (NAA) induction, the E3 ubiquitin ligase complex SCF (Skp1, Cul1, F-box protein TIR1) dimerizes with the mAID-GFP nanobody, resulting in ubiquitination and degradation of GFP-tagged proteins by the proteasome. B) Validation of Pmm2-GFP degradation via Western blot analysis of stage 23 *pmm2-GFP* embryos comparing uninjected to degron injected and induced (50 µM NAA) at 6 hours post fertilization. Gapdh was used as loading control. C) Time-lapse imaging of uninjected homozygous *pmm2-GFP* embryos (n = 7) and degron-injected ones, in ERM (n = 8) or induced with 50 µM NAA (n = 25). D) Quantification of mean GFP fluorescence following baseline correction. Mean and sd are shown. Scale bar: 500 µm. hpi, hours post induction.

To validate the functionality of the degron system in Pmm2-GFP medaka fish, the *TIR1* and *mAID-nanobody* mRNAs of the degron components were co-injected with *mCherry* mRNA as injection tracer into homozygous *pmm2-GFP* embryos at the 1-cell stage. A western blot analysis performed at stage 23 confirmed robust reduction of Pmm2-GFP protein (Fig. 2B). To analyze the kinetics of Pmm2-GFP degradation, the batch of injected embryos was separated and incubated with auxin or kept in embryo rearing medium (ERM) at 6 hours post fertlization (hpf). GFP fluorescence was assesed continuously for 20 hours (Fig. 2C). In uninjected *pmm2-GFP* controls (n = 7), green fluorescence was continuously visible with a slight decrease over time (Fig. 2C-D, green curve). In the group induced with 50 µM NAA (n = 25), the GFP expression quickly decreased (Fig. 2C-D, orange curve; Fig. S3). Degradation of GFP signals also occurred in the yolk of degron-injected and homozygous *pmm2-GFP* individuals which highlights the acute degradation even of maternally provided Pmm2-GFP. Surprisingly, the GFP fluorescence in the degron-system injected but non-induced control group (n = 8) dropped as well (Fig. 2C-D, blue curve; Fig. S3), indicating leakiness of the employed TIR1/mAID system in medaka, i.e. degradation in the absence of auxin.

Although reduction increased by incubation with NAA (Fig 2C-D; Fig. S3), this confirmed that the degron system even without NAA induction harbors a strong basal egradation activity. Due to the leakiness, the TIR1/mAID system was not suitable for use as an acute knockdown application.

### Efficient inducible Pmm2-GFP depletion by the improved TIR1(F74G)/mAID-GFP-nanobody degron system

To overcome leakiness of the TIR1 and mAID/GFP-nanobody degron system, we exchanged the TIR1 F-Box protein with a recently reported variant (F74G), mitigating basal degradation of mAID-tagged proteins (Fig. 3A) (Yesbolatova et al., 2020). The F74G mutation enables the TIR1 protein to bind the auxin analog 5-phenyl-indole-3-acetic acid (5-Ph-IAA) instead of the canonical NAA. Since 5-Ph-IAA can be used at very low concentrations, it minimizes potential toxic effects of this compound. A toxicity screen with varying concentrations of 5-Ph-IAA indeed demonstrated normal embryonic development in medaka fish embryos (Fig. S2).

**Figure 3.**
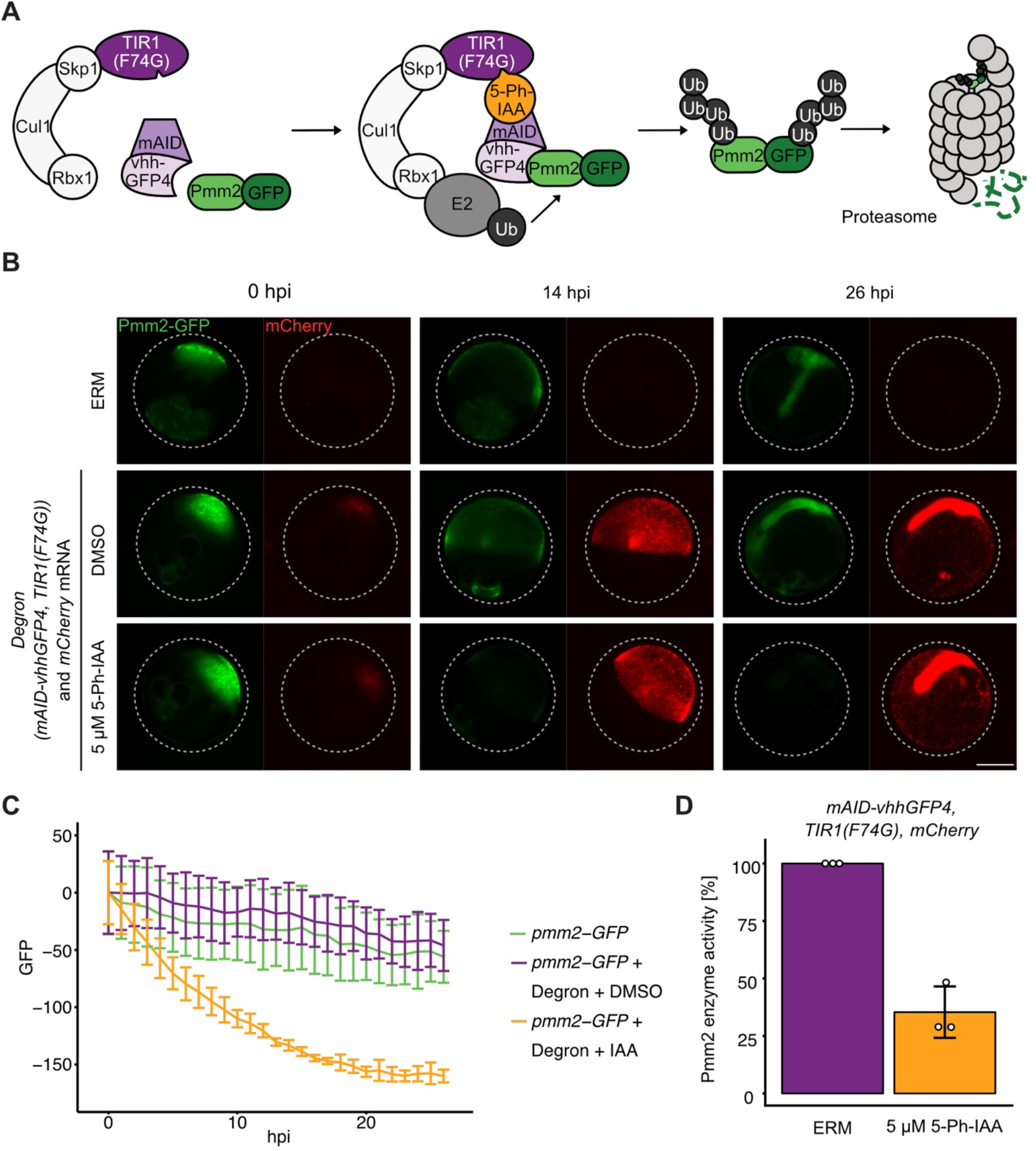
Combination of TIR1(F74G) with mAID/GFP nanobody as degron system enables acute and inducible knockdown of Pmm2-GFP in medaka. A) Schematic representation of the TIR1(F74G) variant combined with the mAID/GFP-nanobody (vhhGFP4) based degron (Daniel et al., 2018; Yesbolatova et al., 2020) to induce degradation of Pmm2-GFP in an auxin-analog (5-Ph-IAA) inducible manner in medaka. B) Time-lapse imaging of uninjected *pmm2-GFP*, degron-injected *pmm2-GFP* embryos incubated in ERM or 5 µM 5-Ph-IAA. C) Quantification of mean GFP fluorescence following baseline correction over 26 hpi. Mean ± sd of triplicates shown, uninjected *pmm2-gfp* control (n total = 15) *degron/mCherry* injected and induced (n total = 65), non-induced embryos (n total = 61). D) Pmm2 enzyme activity assay comparing control (injected, non-induced; purple) and degron injected and induced (5 µM 5-Ph-IAA) embryos (orange). Scale bar: 500 µm. hpi, hour post induction; 5-Ph-IAA, auxin analog Taken together, we show that a mRNA-based systemic degron-induced knockdown setup offers temporal control of protein abundance immediately upon induction, reducing zygotic as well as maternal Pmm2-GFP in medaka. The kinetics of the system revealed fast depletion of Pmm2-GFP with a strong ubiquitous protein degradation and 6.9 hours half-life after induction at non-toxic concentration of 5-Ph-IAA. Especially in the context of disease modeling that aims for early developmental stages or rapid protein turnover, the system is well suited to model the reduction, severe impairment or loss of protein function associated with disease progression. This consequently allows acute modulation of effector enzymes to mimic pathologic levels reported from patients.

To test the specificity and efficiency of TIR1(F74G) to bind the mAID-GFP-nanobody in the presence of 5-Ph-IAA, degron components (*TIR1(F74G)* and *mAID-GFP-nanobody*) were injected as mRNA into 1-cell stage embryos of the homozygous *pmm2-GFP* line along with the injection tracer *mCherry*. The injected embryos were split into treated (5 µM 5-Ph-IAA; n total = 65) and untreated (n total = 61) groups.

Green (Pmm2-GFP) fluorescence was acquired by time-lapse imaging over 26 hpi (Fig. 3B). In the uninjected *pmm2-GFP* control groups (n total = 15), green fluorescence was continuously visible with a slight decrease over time (Fig. 3B-C, green curve; Fig. S4). In the degron injected but non-induced groups, the GFP fluorescence followed the same trend as the uninjected control, indicating no obvious basal degradation of the TIR1(F74G)/mAID-vhhGFP4 system (Fig. 3B-C, purple curve; Fig. S4). An immediate decrease in GFP fluorescence was observed in the groups induced with 5 µM 5-Ph-IAA, reaching a plateau about 20 hours post induction (Fig. 3B-C, Fig. S4). Based on fitted curve analysis, Pmm2-GFP was degraded with a half-life of approximately 6.9 hours (Fig. S5). Compared to degron injected and non-induced Pmm2-GFP control embryos (n total = 85 of three independent replicates), a biochemical Pmm2 enzyme activity assay at stage 23 confirmed the reduction of net Pmm2 activity to below 50 % in degron injected and induced (5 µM 5-Ph-IAA) embryos (n total = 85 of three independent replicates) (Fig. 3D).

The combined use of TIR1(F74G) and mAID-GFP-nanobody thus allowed acute knockdown of proteins endogenously tagged with GFP in a developing medaka embryo. In the case of Pmm2-GFP, this system reduced the net enzyme activity of Pmm2 to pathologic levels reported from PMM2-CDG patients.

For Pmm2-GFP, this opens the possibility to generate a hypo-glycosylation environment at early stages of development, challenging to reach with classic genetic methods which rely on *pmm2* mutant alleles. Homozygous animals often cannot be raised to maturity due to their severe pathology. Heterozygous breeds in contrast are usually not haploinsufficient and enzyme levels exceed the phenotypic threshold. Offspring of heterozygous mothers in turn do not allow to study the early onset of CDG-phenotypes, as Pmm2 is maternally contributed, mitigating the early embryonic phenotype.

Comparing previous reports of different TIR1/AID systems and its genetic variants highlight that there is not a single solution generally applicable across different model systems and that TIR1/AID is frequently reported to show basal activity (Schiksnis et al., 2020). The system was improved by mutating the phenylalanine in position 74 in the TIR1 protein, resulting in different efficiency depending on the organism used (*Drosophila melanogaster, Caenorhabditis elegans, Oryzias latipes, Schizosaccharomyces pombe)* (Goldner et al., 2023; Hills-Muckey et al., 2022; Kiyomitsu et al., 2024; Zhang et al., 2022). In yeast, a phenylalanine to alanine change is reported to be more effective compared to other mutations (Ogawa et al., 2023). In cell culture systems, comparable levels of protein degradation were observed for both F74A and G variants. However, F74A showed higher activity with IAA, preferring the usage of F74G variant in mouse (Yesbolatova et al., 2020). Further, there is a clear improvement in using the TIR1-variant sensitive to auxin analogs (5-Ad-IAA; 5-Ph-IAA) versus the native plant hormone when it comes to toxicity effects of the amounts required for the induction of protein knockdown (Nishimura et al., 2020; Yesbolatova et al., 2020). In the end it is advisable to test reported alternatives like the TIR1(F74G)/mAID-GFP-nanobody system as it may vary in basal activity or effectivity in the used species or cell system.

For developmentally relevant time windows requiring acute protein degradation, the mRNA-based approach provides many advantages. It gives immediate access to any endogenously GFP tagged target protein and at the same time it overcomes the need for universal, ubiquitous promoters active during early embryogenesis. The degradation efficiency can immediately be adjusted (by the amount of injected mRNA) to the levels of expression of the targeted protein of interest, a requirement that is hard to meet in fixed and stable transgenic settings. The system presented thus offers a fast and immediate approach for highly efficient acute protein degradation of developmentally relevant proteins and provides an important complementation to genetic targeted inactivation, in particular in settings with a strong maternal contribution.

## Material and Methods

### Fish maintenance

Medaka (*Oryzias latipes*) stocks were maintained (fish husbandry, permit number 35– 9185.64/BH Wittbrodt) and experiments (permit number 35–9185.81/G-271/20 Wittbrodt) were performed in accordance with local animal welfare standards (Tierschutzgesetz §11, Abs. 1, Nr. 1) and European Union animal welfare guidelines (Bert et al., 2016). Fish were maintained in closed stocks and constant recirculating systems at 28 °C on a 14 h light/10 h dark cycle. The fish facility is under the supervision of the local representative of the animal welfare agency.

### Line generation

The endogenously tagged *pmm2-GFP* lines was generated as described in Gutierrez-Triana et al. (2018). In short: The *pmm2* template plasmid containing GFP sequence flanking 5’ and 3’homology flanks (HFs) was created via GoldenGATE cloning described in Kirchmaier et al. (2013). Biotinylated primers were used to amplify a modified PCR doner. To minimize interference of Pmm2 stability and folding capacity, a flexible GS linker composed of 21 amino acids upstream of GFP (Sabourin et al., 2007) was used. Wild-type Cab embryos were injected with a sgRNA targeting the *pmm2* locus at the C-terminus (5’-TCTTCTTCTGCTGAAGCTAC[TGG]-3’ and 5’-TCTTCTGCTGAAGCTACTGG[AGG]-3’), *heiCas9* mRNA, and *mCherry* mRNA as injection tracer. Positively screened embryos were raised to adulthood and outcrossed to wild-type medakas. The resulting generation was screened for ubiquitous GFP expression. Founder fish were outcrossed to raise stable heterozygous individuals, following generation of a fertile homozygous *pmm2-GFP* line. Single integration was verified by Southern blot, PCR genotyping, and Western blot (Fig. S1).

### Genotyping – gDNA and cDNA analysis

Genomic DNA (gDNA) was extracted from fin clips using fin-clip buffer (0.4 M Tris-HCl pH 8.0, 5 mM EDTA pH 8.0, 0.15 M NaCl, 0.1 % SDS in distilled water with 1 mg/ml Proteinase K (Roche, 20 mg/ml)) over night at 60 °C. After incubation, 2 Volumes of nuclease free water was added and lysates were incubated at 95 °C for 20 min. For gDNA extraction from embryos, up to 5 individuals were ground with plastic pestles in 100 μl Fin-Clip buffer and 5µl Proteinase K (20 mg/ml), incubated for 1 h at 60 °C. After addition of 200 µl nuclease free water, lysates were incubated at 95 °C for 20 min. Samples were stored at 4 °C. Genotyping was performed using 1 μl of lysate in a 50 μl PCR reaction with 1x Q5 reaction buffer (New England Biolabs, NEB), 200 µM dNTPs (Sigma), 200 µM per primer (Eurofins Genomics) and 0.012 U/µl Q5 polymerase (NEB). *Pmm2* locus primers were used: *Pmm2-fwd* 5’-TGGACGACACAGATGATGCT and *Pmm2-rev* 5’-CTCACAGACTGACCCTCACC. PCRs were run for 30 cycles (30 sec, 98 °C denaturation; 30 sec, 67 °C annealing, 90 sec, 72°C extension). Amplicons were size fractioned by gel electrophoresis on 1 % agarose gel in 1x TAE (40mM Tris, 20mM acetic acid, 1mM EDTA) and sent for Sanger sequencing (Eurofins Genomics) after gel extraction and purification (monarch kit, NEB).

For RT-PCT, total RNA was extracted from five *pmm2-GFP* and wild-type hatchlings by lysis in TRIzol (Ambion) and purification with the Direct-zol RNA MicroPrep Kit (Zymo Research) according to the manufacturers protocol. The complementary DNA (cDNA) was prepared by reverse transcription using Revert Aid Kit (Thermo Fisher Scientific). Primers binding to exon 2 of *Pmm2* (5’-AAACGCAGAGGCTCAGGACTCG) and the 3’UTR (5’-TGAGGTCACATCCCGTGTTG) were used for a 30 cycles PCR as described above (68 °C annealing, 2 min 72 °C extension). PCR products were run on a 1.5 % agarose gel in 1x TAE and sent for sanger sequencing after filter-column purification from the gel (Monarch, NEB).

### Protein extraction, quantification and immunoblot analysis for Pmm2-GFP line validation

For validation of the Pmm2-GFP line, protein lysates for immunoblot analysis were prepared from hatchlings homogenized in 1x PBS (137 mM NaCl, 2.7 mM KCl, 1.44 g/L Na2HPO4, 240 mg/L KH2PO4) with 1 % TritonX with 1x cOmplete EDTA– free Protease Inhibitor Cocktail (Roche). After incubation for 20 min on the rotator in the cold room, samples were centrifuged at 12,000 g at 4 °C for 10 min. The supernatant was transferred to a low protein binding Eppendorf tube. The concentrations of the lysates were determined with BCA protein-assay kit (23225, Thermo Fisher Scientific). 25 µg of protein lysate was boiled in 2.5x Laemmli sample buffer (157.5 mM Tris-HCl [pH 6.8], 5% SDS, 25% glycerol, 12.5% 2-mercaptoethanol, 0.08% Bromophenol Blue) for 10 min at 95 °C and stored at −80 °C. The samples were run on a self-made SDS-PAGE with 10 % separation gel and 4 % stacking gel at 60 V for the first 30 min and at 120 V afterwards. Transfer of the proteins to a PVDF membrane (IPVH00010, Millipore Immobilon-P) was done at 350 mA for 1 h at 4 °C. Membranes were blocked in 5 % Skim Milk (Sigma-Aldrich) in 1xTBST (50 mM Tris-HCl [pH 7.5], 150 mM NaCl, 0.1% (v/v) Tween 20) for 1 – 2 h at room temperature (RT). Blots were incubated with primary antibody in blocking solution over night at 4 °C. Antibodies used: anti-GFP (A11122, Invitrogen, 1:500), anti-PMM2 (Proteintech, 10666-1-AP, 1:500), anti-GAPDH (Cell singling 14C10C21188, 1:1000). Blots were washed 4x 10 min with 1x TBST and incubated afterwards with corresponding secondary antibody (Goat anti-rabbit – Santa Cruz (sc-2005), 1:10000) diluted in 1x TBST for 1 h at RT. Blots were washed with 1x TBST 4x 10 min. Proteins were visualized with ECL Substrate (Pierce ECL Western Blotting Substrate) on a Gel Doc Imager (INTAS, Göttingen).

### Southern blot

To validate single integration of *GFP* into the *pmm2* locus, Southern blot was performed on gDNA extracted from homozygous *pmm2-GFP* adult heads. gDNA was isolated by ethanol precipitation after phenol-chloroform extraction and resuspended in 1x TE buffer (10 mM Tris HCl pH 8.0, 1 mM EDTA pH 8.0). 10 µg *pmm2-GFP* gDNA and 200 pg of *GFP* control plasmid were digested with 10 U of BspHI (NEB) in combination with 20 U of HindIII (NEB) or 20 U of BsaI HF (NEB) at 37 °C over night. The digested fragments were size fractionated by electrophoresis on a 0.8 % agarose gel in 1x TAE buffer. On the gel, DNA was depurinated in 0.25 N HCl for 30 min at RT, rinsed with H_2_O, denatured in 0.5 N NaOH, 1.5 M NaCl solution for 30 min at RT and neutralized in 0.5 M Tris HCl, 1.5 M NaCl, pH 7.2 before it was transferred overnight at room temperature onto a Hybond-N+ membrane (Sigma-Aldrich) by capillary transfer. The membrane was washed with 50 mM NaPi for 5 min at RT, DNA was UV crosslinked and pre-hybridized in Church hybridization buffer (0.5 M NaPi, 7% SDS, 1 mM EDTA pH 8.0) at 65 °C for 30 min. Probe synthesis was performed with PCR DIG Probe Synthesis Kit (Roche) according to the manufacturer’s protocols (primers used: GFPprobe-fwd 5’-GTGAGCAAGGGCGAGGAGCT, GFPprobe-rev 5’-TTACTTGTACAGCTCGTCCATG) and PCR conditions: initial denaturation at 95 °C for 2 min, 35 cycles of 95 °C 30 s, 60 °C 30 s, 72 °C 40 s and final extension at 72 °C 7 min. The probe comprised a 700 bp fragment containing *GFP* sequence. The probe was denatured in hybridization buffer for 10 min at 95 °C and hybridized with the membrane overnight at 65 °C. The membrane was washed with pre-heated (65°C) Church washing buffer (40 mM NaPi, 1% SDS) at 65 °C for 10 min, continued at RT for 10 min and washed with 1x DIG1 % and 0.3 % Tween (Sigma-Aldrich) for 5 min at RT. The membrane was blocked in 1 % w/v blocking reagent (Roche) in 1x DIG1 solution at RT for at least 30 min. The membrane was incubated with 1:10,000 anti-digoxigenin-AP Fab fragments (Roche) for 30 min at RT in 1 % w/v blocking reagent (Roche) in 1x DIG1 solution. Two washing steps with 1x DIG1 % and 0.3 % Tween were performed for 20 min at RT followed by a 5 min washing step in 1x DIG3 (0.1 M Tris pH 9.5, 0.1 M NaCl) at RT. Detection was performed using 6 µl/ml CDP star (Roche) and chemiluminescence was acquired on an Intas Imager (INTAS, Göttingen).

### Auxin toxicity test

For the auxin toxicity test a 0.1 M stock solution was prepared by dissolving NAA powder (NAA, sodium 1-naphtaleneacetate, Santa Cruz Biotechnology) in 1x ERM. Working concentrations were further diluted with ERM from the stock solution. Wild-type Cab embryos from adult crosses were collected and kept until stage 10 (~6 hpf) in ERM. Next, embryos were induced with 0.1 mM, 0.5 mM, 1 mM, 5 mM and 10 mM NAA and kept at 26 °C. Auxin solution was refreshed every 24 hours. The resulting phenotypes were assessed by imaging every day until 4 dpf under a stereomicroscope with an attached Nikon DXM1200 digital camera.

For the auxin analog 5-Ph-IAA toxicity test, the powder (MedChemExpress, # HY-134653) was dissolved in DMSO and a 10 mM stock solution was prepared in 1x PBS. Working concentrations were further diluted with ERM from the stock solution. Wild-type Cab embryos from adult crosses were collected and kept until stage 10 (~6 hpf) in ERM. Next, embryos were induced with 1 µM, 2.5 µM, 5 µM and 10 µM 5-Ph-IAA and kept at 26 °C. Auxin solution was refreshed every 48 hours. The resulting phenotypes were assessed by imaging every day until hatch under a stereomicroscope with the attached Nikon camera.

### Degron system plasmids and mRNAs synthesis

The degron system mRNAs were synthesized from constructs (Daniel et al., 2018) cloned into pCS2+ plasmids. pCDNA5FRT/TO_HA-mAID-nanobody was a gift from Joerg Mansfeld (Addgene plasmid # 117713; http://n2t.net/addgene:117713; RRID:Addgene_117713) and pCS2+_Flag-myc-NES-Tir1 was as well a gift from Joerg Mansfeld (Addgene plasmid # 117717; http://n2t.net/addgene:117717; RRID:Addgene_117717). TIR1F74G point mutation (Yesbolatova et al., 2020) was introduced via Q5 mutagenesis.

For *in vitro* transcription, 5 µg of the plasmids were linearized O/N at 37 °C and the digests were purified with QIAquick PCR Purification Kit (Qiagen). *In vitro* transcriptions of mRNAs were prepared with the mMESSAGE mMACHINE SP6 Transcription Kit (Invitrogen) and purified with the RNeasy Mini Kit (Qiagen), according to manufacturers’ protocols. The mRNA quality was assessed through a RNA agarose test gel.

### Microinjections

For the validation of degron systems, microinjections were performed in homozygous *pmm2-GFP* embryos at the one-cell stage. Freshly fertilized eggs were injected with respective injection mixes. After injections, embryos were transferred into fresh embryo-rearing medium (1× ERM: 17 mM NaCl, 40 mM KCl, 0.27 mM CaCl2•2H2O, 0.66 mM MgSO4•7H2O and 17 mM HEPES) and incubated at 26 °C. 6 hours after fertilization embryos were induced with auxin or the auxin analog 5-Ph-IAA. Prior to induction, dead and mCherry negative embryos were discarded.

### Time-lapse acquisition and quantification of Pmm2-GFP degradation

Time-lapse images of degron-injected *pmm2-GFP* embryos were acquired on a ACQUIFER imaging Machine (ACQUIFER Imaging GmbH, Heidelberg, Germany). Embryos were placed in a 96-well plate with 200 µl media and inserted into the plate holder of the imaging machine. Imaging was performed at 26 °C for 20 or 26 hours. A set of 10 z-slices (10 µm each slice) was acquired in bright field, 561 nm and 470 nm fluorescence channels with 2x NA 0.06 objective (Nikon, Düsseldorf, Germany) every or every 2 hours.

For analysis of the degron kinetics a self-written ImageJ macro was used. In brief, for each timepoint, a maximum projection of the z-stack was calculated for both fluorescence images. Next, the maximum-projection images were cropped by a rectangular mask in which the embryos lie in the middle to exclude background. The fluorescence intensity from each embryo for each time-point was used for analysis of degradation kinetics of GFP. The mean GFP was baseline normalized, i.e. subtracting the initial time zero value from every point in the curve.

The plots were visualized with R.

### Validation of degradation via western blot

Pmm2-GFP protein degradation was validated via western blot. For sample preparation, stage 23 embryos were gently rolled on sandpaper to remove the outer hair and treated afterwards in glass vials with hatching enzyme for ~45 min at 28 °C. Next, embryos were washed 4x with ERM and transferred into a glass dish filled with ERM. Remnants of the chorion were removed, and the yolk was teared open with forceps. Samples were centrifuged for 2 min at 956 g at 4 °C. Afterwards, the supernatant was removed and the samples were smashed in lysis buffer with a plastic pestle. The subsequent procedure for protein lysates and western blot analysis, is like described before for preparation of protein lysates from hatchlings in “Protein extraction, quantification and immunoblot analysis for *pmm2-GFP* line validation” section.

### Pmm2 enzyme activity assay

Protein lysates were extracted from stage 23 degron injected embryos and the procedure was the same as describes above in the “Validation of degradation via western blot” section. After the last centrifugation step, the samples were snap-frozen in liquid nitrogen and stored at −80 °C. The sample pellets were resuspended in lysis buffer which contained protease inhibitors without detergent. Measurement of the Pmm2 activity followed the procedure of Körner et al., 1998. In short, 30 μg of protein were mixed with NADP, MgCl_2_, glucose-1-phosphate, mannose-6 phosphate isomerase (MPI) and phosphoglucose isomerase (PGI). In order to use Pmm2 as a rate-limiting step of the reaction, both enzymes, MPI and PGI, were supplemented in excessive amount to oversaturate them. The conversion of NADP to NADPH was measured at 340 nm for 2 hours. Samples were measured in triplicates.

## Acknowledgement

Alicia Pérez Saturnino cloned the Pmm2-GFP plasmid for the production of modified PCR. Simran Panda screened Pmm2-GFP fish. Elena Tonin performed the cloning of TIR1 and AID-nanobody into PCS2+ vector for *in vitro* mRNA synthesis. Virginia Geiger performed Pmm2 enzyme assay. Johanna Rasch performed the cDNA preparation.

## Author contributions

Conceptualization: K.P., J.W., T.T.; Data curation: K.P., S.J., E.S.S.; Formal analysis: K.P., S.J., E.S.S., T.T., Funding acquisition: T.T., J.W., C.T.; Investigation: K.P., S.J., E.S.S.; Methodology: K.P., S.J., T.T., C.T.; Project administration: J.W., T.T.; Resources: J.W., C.T.; Supervision: J.W., T.T.; Visualization: K.P.; Writing – original draft: K.P., T.T., J.W. with input from S.J., E.S.S., C.T.

## Funding

This research was supported by grants from the Deutsche Forschungsgemeinschaft (DFG) through FOR2509 project 10 to J.W. (WI1824/9-1) and to T.T. (TH1992/1-2) as well as to C.T. project 9 (TH1461/7-2), and the European Research Council (ERC-SyG H2020, 810172 to J.W.), and the Excellence Cluster ‘3D Matter Made to Order’ (3DMM2O, EXC 2082/1 Wittbrodt C3) funded through the German Excellence Strategy via Deutsche Forschungsgemeinschaft (DFG) to J.W.

## Data and resource availability

All relevant data can be found within this article and the supplementary information

## Competing interests

The authors declare no competing or financial interests.

## Notes

### Competing Interest Statement

The authors have declared no competing interest.

## References

Caussinus, E., & Affolter, M. (2016). deGradFP: A System to Knockdown GFP-Tagged Proteins (pp. 177–187). 10.1007/978-1-4939-6371-3_9

Caussinus, E., Kanca, O., & Affolter, M. (2011). Fluorescent fusion protein knockout mediated by anti-GFP nanobody. Nature Structural & Molecular Biology, 19(1), 117–121. 10.1038/nsmb.2180

Chan, B., Clasquin, M., Smolen, G. A., Histen, G., Powe, J., Chen, Y., Lin, Z., Lu, C., Liu, Y., Cang, Y., Yan, Z., Xia, Y., Thompson, R., Singleton, C., Dorsch, M., Silverman, L., Su, S.-S. M., Freeze, H. H., & Jin, S. (2016). A mouse model of a human congenital disorder of glycosylation caused by loss of PMM2. Human Molecular Genetics, 25(11), 2182–2193. 10.1093/hmg/ddw085

Cline, A., Gao, N., Flanagan-Steet, H., Sharma, V., Rosa, S., Sonon, R., Azadi, P., Sadler, K. C., Freeze, H. H., Lehrman, M. A., & Steet, R. (2012). A zebrafish model of PMM2-CDG reveals altered neurogenesis and a substrate-accumulation mechanism for N-linked glycosylation deficiency. Molecular Biology of the Cell, 23(21), 4175–4187. 10.1091/mbc.e12-05-0411

Daniel, K., Icha, J., Horenburg, C., Müller, D., Norden, C., & Mansfeld, J. (2018). Conditional control of fluorescent protein degradation by an auxin-dependent nanobody. Nature Communications, 9(1), 3297. 10.1038/s41467-018-05855-5

Doering, L., Cornean, A., Thumberger, T., Benjaminsen, J., Wittbrodt, B., Kellner, T., Hammouda, O. T., Gorenflo, M., Wittbrodt, J., & Gierten, J. (2023). CRISPR-based knockout and base editing confirm the role of MYRF in heart development and congenital heart disease. Disease Models & Mechanisms, 16(8). 10.1242/dmm.049811

Goldner, A. N., Fessehaye, S. M., Rodriguez, N., Mapes, K. A., Osterfield, M., & Doubrovinski, K. (2023). Evidence that tissue recoil in the early Drosophila embryo is a passive not active process. Molecular Biology of the Cell, 34(10). 10.1091/mbc.E22-09-0409

Gücüm, S., Sakson, R., Hoffmann, M., Grote, V., Becker, C., Pakari, K., Beedgen, L., Thiel, C., Rapp, E., Ruppert, T., Thumberger, T., & Wittbrodt, J. (2021). A patient-based medaka alg2 mutant as a model for hypo-N-glycosylation. Development, 148(11). 10.1242/dev.199385

Gutierrez-Triana, J. A., Tavhelidse, T., Thumberger, T., Thomas, I., Wittbrodt, B., Kellner, T., Anlas, K., Tsingos, E., & Wittbrodt, J. (2018). Efficient single-copy HDR by 5’ modified long dsDNA donors. ELife, 7. 10.7554/eLife.39468

Hammouda, O. T., Wu, M. Y., Kaul, V., Gierten, J., Thumberger, T., & Wittbrodt, J. (2021). In vivo identification and validation of novel potential predictors for human cardiovascular diseases. PLOS ONE, 16(12), e0261572. 10.1371/journal.pone.0261572

Hills-Muckey, K., Martinez, M. A. Q., Stec, N., Hebbar, S., Saldanha, J., Medwig-Kinney, T. N., Moore, F. E. Q., Ivanova, M., Morao, A., Ward, J. D., Moss, E. G., Ercan, S., Zinovyeva, A. Y., Matus, D. Q., & Hammell, C. M. (2022). An engineered, orthogonal auxin analog/ At TIR1(F79G) pairing improves both specificity and efficacy of the auxin degradation system in Caenorhabditis elegans. Genetics, 220(2). 10.1093/genetics/iyab174

Himmelreich, N., Kaufmann, L. T., Steinbeisser, H., Körner, C., & Thiel, C. (2015). Lack of phosphomannomutase 2 affects Xenopus laevis morphogenesis and the non-canonical Wnt5a/Ror2 signalling. Journal of Inherited Metabolic Disease, 38(6), 1137–1146. 10.1007/s10545-015-9874-0

Kiyomitsu, A., Nishimura, T., Hwang, S. J., Ansai, S., Kanemaki, M. T., Tanaka, M., & Kiyomitsu, T. (2024). Ran-GTP assembles a specialized spindle structure for accurate chromosome segregation in medaka early embryos. Nature Communications, 15(1), 981. 10.1038/s41467-024-45251-w

Korner, C., Lehle, L., & von Figura, K. (1998). Abnormal synthesis of mannose 1-phosphate derived carbohydrates in carbohydrate-deficient glycoprotein syndrome type I fibroblasts with phosphomannomutase deficiency. Glycobiology, 8(2), 165–171. 10.1093/glycob/8.2.165

Natsume, T., Kiyomitsu, T., Saga, Y., & Kanemaki, M. T. (2016). Rapid Protein Depletion in Human Cells by Auxin-Inducible Degron Tagging with Short Homology Donors. Cell Reports, 15(1), 210–218. 10.1016/J.CELREP.2016.03.001

Ng, B. G., Freeze, H. H., Himmelreich, N., Blau, N., & Ferreira, C. R. (2024). Clinical and biochemical footprints of congenital disorders of glycosylation: Proposed nosology. Molecular Genetics and Metabolism, 142(1), 108476. 10.1016/j.ymgme.2024.108476

Nishimura, K., Fukagawa, T., Takisawa, H., Kakimoto, T., & Kanemaki, M. (2009). An auxin-based degron system for the rapid depletion of proteins in nonplant cells. Nature Methods, 6(12), 917–922. 10.1038/nmeth.1401

Nishimura, K., Yamada, R., Hagihara, S., Iwasaki, R., Uchida, N., Kamura, T., Takahashi, K., Torii, K. U., & Fukagawa, T. (2020). A super-sensitive auxin-inducible degron system with an engineered auxin-TIR1 pair. Nucleic Acids Research, 48(18), e108–e108. 10.1093/nar/gkaa748

Ogawa, Y., Nishimura, K., Obara, K., & Kamura, T. (2023). Development of AlissAID system targeting GFP or mCherry fusion protein. PLOS Genetics, 19(6), e1010731. 10.1371/journal.pgen.1010731

Pajusalu, S., Vals, M.-A., Serrano, M., Witters, P., Cechova, A., Honzik, T., Edmondson, A. C., Ficicioglu, C., Barone, R., De Lonlay, P., Bérat, C.-M., Vuillaumier-Barrot, S., Lam, C., Patterson, M. C., Janssen, M. C. H., Martins, E., Quelhas, D., Sykut-Cegielska, J., Mousa, J., … Õunap, K. (2024). Genotype/Phenotype Relationship: Lessons From 137 Patients With PMM2-CDG. Human Mutation, 2024(1). 10.1155/2024/8813121

Parkinson, W. M., Dookwah, M., Dear, M. L., Gatto, C. L., Aoki, K., Tiemeyer, M., & Broadie, K. (2016a). Neurological roles for phosphomannomutase type 2 in a new Drosophila congenital disorder of glycosylation disease model. Disease Models & Mechanisms. 10.1242/dmm.022939

Parkinson, W. M., Dookwah, M., Dear, M. L., Gatto, C. L., Aoki, K., Tiemeyer, M., & Broadie, K. (2016b). Synaptic roles for phosphomannomutase type 2 in a new Drosophila congenital disorder of glycosylation disease model. Disease Models & Mechanisms, 9(5), 513–527. 10.1242/dmm.022939

Schiksnis, E., Nicholson, A., Modena, M., Pule, M., Arribere, J., & Pasquinelli, A. (2020). Auxin-independent depletion of degron-tagged proteins by TIR1. MicroPublication Biology, 2020. 10.17912/micropub.biology.000213

Schneider, A., Thiel, C., Rindermann, J., DeRossi, C., Popovici, D., Hoffmann, G. F., Gröne, H.-J., & Körner, C. (2012). Successful prenatal mannose treatment for congenital disorder of glycosylation-Ia in mice. Nature Medicine, 18(1), 71–73. 10.1038/nm.2548

Sharma, V., Ichikawa, M., & Freeze, H. H. (2014). Mannose metabolism: More than meets the eye. Biochemical and Biophysical Research Communications, 453(2), 220–228. 10.1016/j.bbrc.2014.06.021

Thiel, C., Lübke, T., Matthijs, G., von Figura, K., & Körner, C. (2006). Targeted Disruption of the Mouse Phosphomannomutase 2 Gene Causes Early Embryonic Lethality. Molecular and Cellular Biology, 26(15), 5615–5620. 10.1128/MCB.02391-05

Wittbrodt, J., Shima, A., & Schartl, M. (2002). Medaka — a model organism from the far east. Nature Reviews Genetics, 3(1), 53–64. 10.1038/nrg704

Yesbolatova, A., Saito, Y., Kitamoto, N., Makino-Itou, H., Ajima, R., Nakano, R., Nakaoka, H., Fukui, K., Gamo, K., Tominari, Y., Takeuchi, H., Saga, Y., Hayashi, K., & Kanemaki, M. T. (2020). The auxin-inducible degron 2 technology provides sharp degradation control in yeast, mammalian cells, and mice. Nature Communications, 11(1), 5701. 10.1038/s41467-020-19532-z

Zhang, X.-R., Zhao, L., Suo, F., Gao, Y., Wu, Q., Qi, X., & Du, L.-L. (2022). An improved auxin-inducible degron system for fission yeast. G3 Genes|Genomes|Genetics, 12(1). 10.1093/g3journal/jkab393

Zhong, M., & Lai, K. (2025). AAV-based gene replacement therapy prevents and halts manifestation of abnormal neurological phenotypes in a novel mouse model of PMM2-CDG. Gene Therapy. 10.1038/s41434-025-00525-w

