## Supplemental Figures for "Establishing an auxin-inducible GFP nanobody-based acute protein knockdown system to mimic hypomorphic mutations during early medaka embryogenesis"

### Supplementary Figures

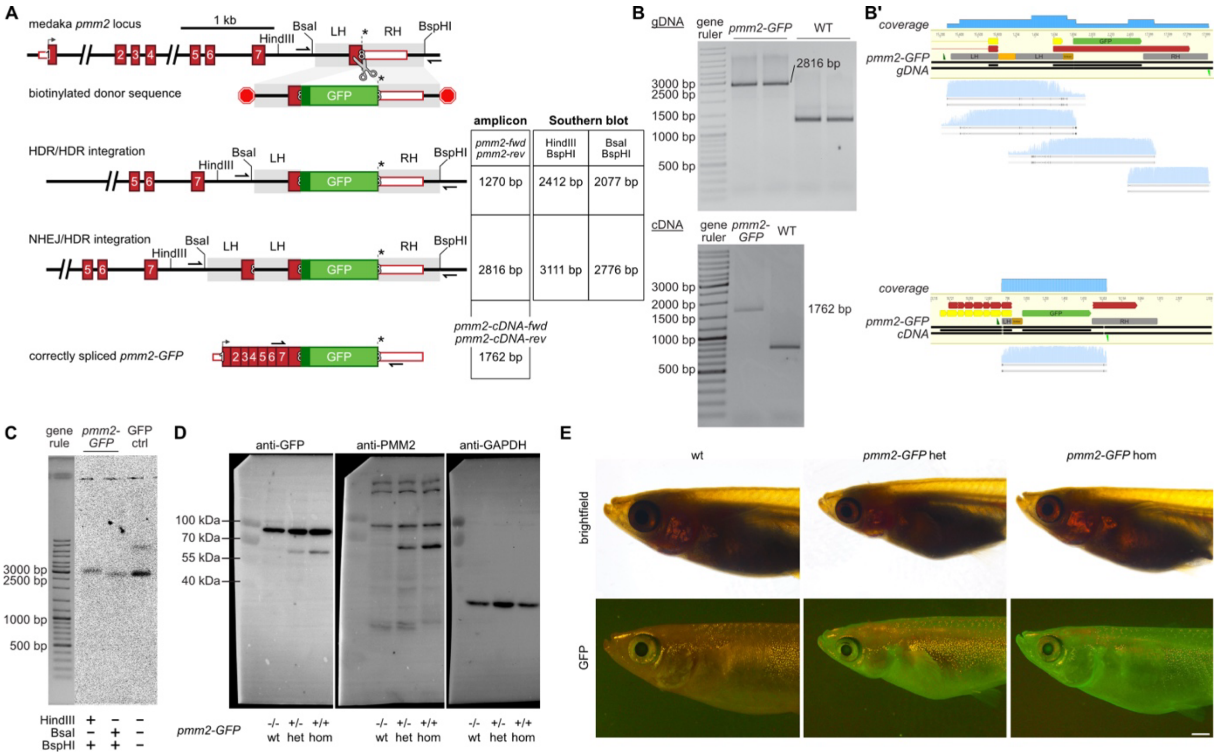


**Supplementary Figure 1: Validation of *pmm2-GFP* tagged line.** A) Possible outcomes of GFP integration into Pmm2 locus, their sizes after locus-locus PCR and restriction enzyme cutting sites for southern blot analysis. B) gDNA and cDNA sequencing of homozygous *pmm2-GFP* hatchlings. Locus-locus PCR on gDNA indicates NHEJ event. Sanger sequencing confirms 5' NEHJ and 3'HDR integration (red – mRNA, yellow - CDS). C) Southern blot analysis of homozygous *pmm2-GFP* adults confirm single GFP integration. Single band hybridization signals were detected when cutting outside and within the 3'HF with two sets of enzymes (BsaI/BspHI and HindIII/BspI). D) Whole western blots from analysis of lysates from *pmm2*/*pmm2* wild-type, pmm2/*pmm2-GFP* heterozygous and *pmm2-GPF/pmm2-GFP* homozygous hatchlings from Fig. 1. Pmm2 and GFP specific antibodies were used and Gapdh was used as loading control. E) Adult *pmm2/pmm2* wild-type, *pmm2/pmm2-GFP* heterozygous and *pmm2-GPF/pmm2-GPF* homozygous medaka fish. Scale bar 1 mm. cDNA, complementary DNA; gDNA, genomic DNA; HDR, homology directed repair, het, heterozygous; hom, homozygous; NHEJ, non-homologous end-joining; wt, wild-type

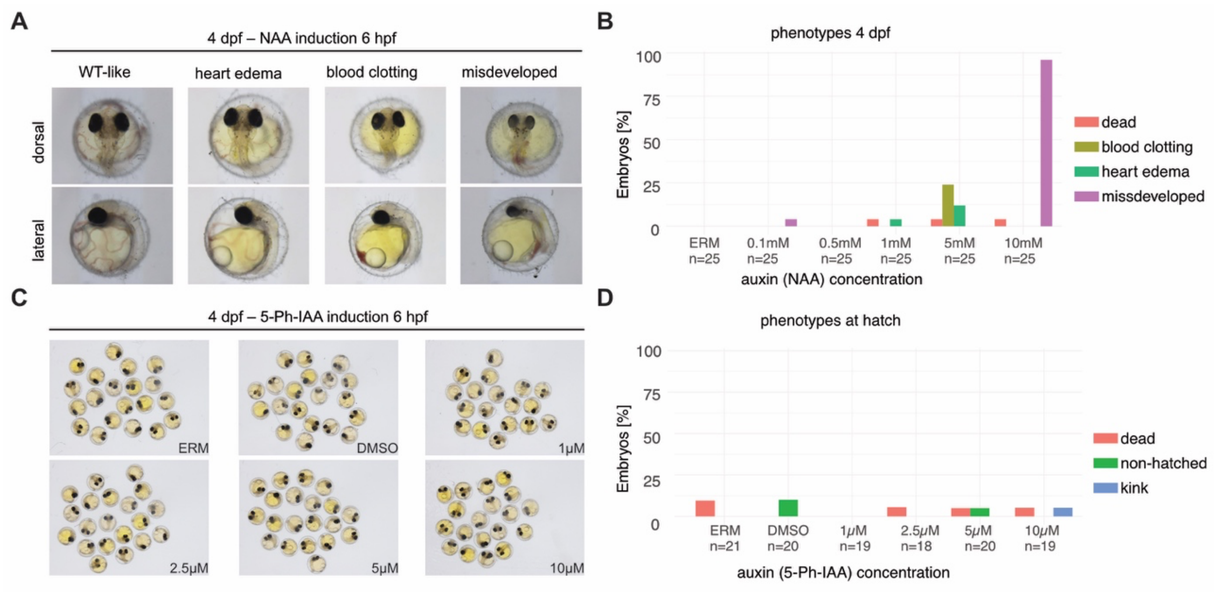


**Supplementary Figure 2 Auxin toxicity in wild-type embryos.** A) Representative NAA induced phenotypes resulting from incubation in auxin solutions 6hpf with different concentrations. B) Quantification of NAA induced phenotypes from an auxin (NAA) toxicity test in wild-type embryos 4 dpf. C) Representative 5-Ph-IAA induced phenotypes resulting from incubation in auxin solutions with different concentrations 6hpf. D) Quantification of 5-Ph-IAA induced phenotypes from toxicity test 6hpf. dpf, days post fertilization; hpf, hours post fertilization

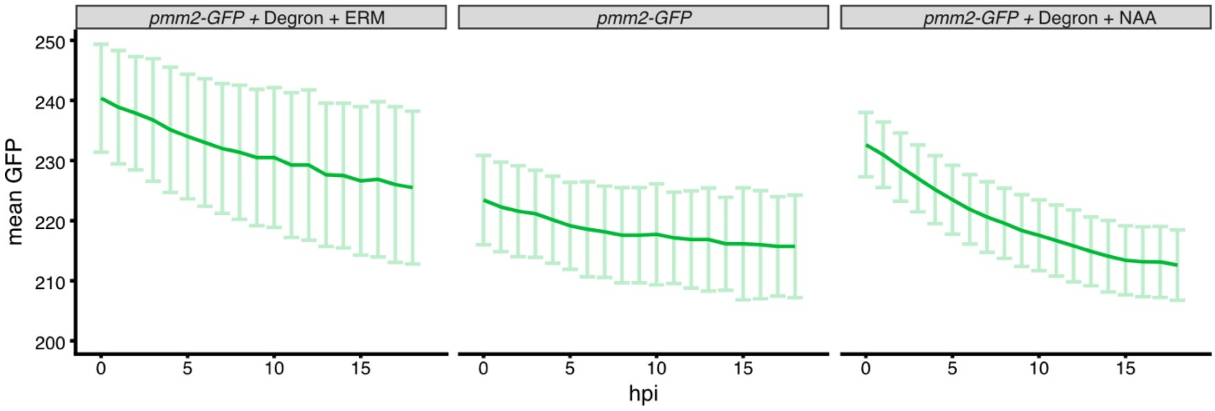


**Supplementary Figure 3 Mean GFP fluorescence acquisition of control and degron injected group.** Temporal fluorescence acquisition in non-injected homozygous pmm2-GFP embryos (n=7) and degron injected control (n=8) and treated group (n=25). Mean values ± sd. n. Rawdata to data shown in Fig. 2D

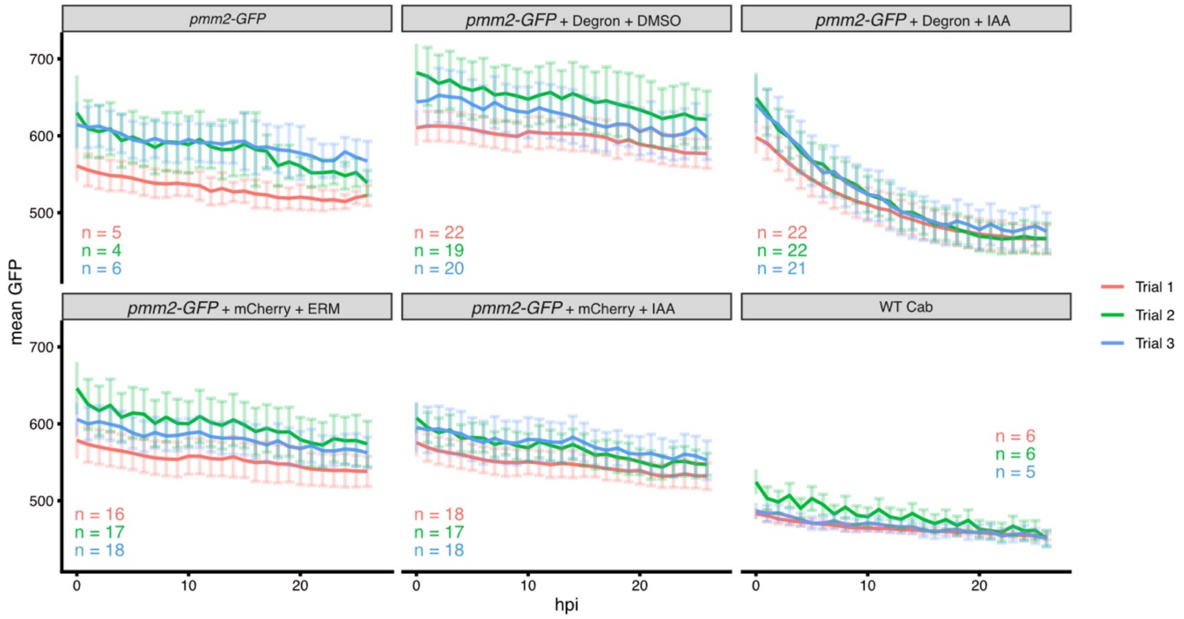


**Supplementary Figure 4 Quantification of mean GFP fluorescence of control and degron injected groups.** Temporal mean GFP fluorescence acquisition over 26 hours in control, degron injected and treated groups. Mean values ± sd given per trial. N numbers provided in figure. Rawdata to triplicates used in Fig. 3C.

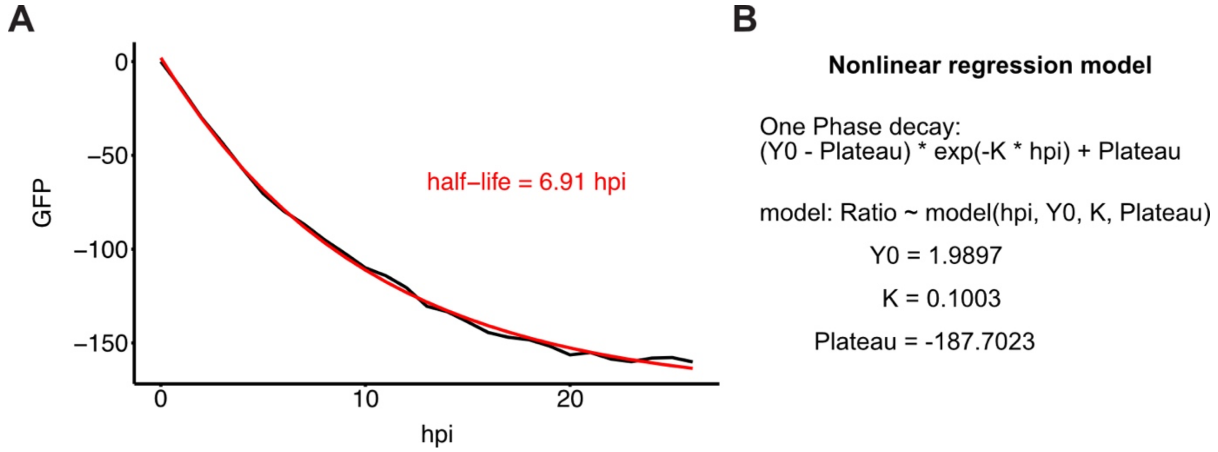


**Supplementary Figure 5 One-phase exponential decay model to determine half-life of Pmm2-GFP degradation.** A) Baseline corrected mean value of GFP fluorescence fitted to a one-phase exponential decay model (Y = (Y0-Plateau)exp(-k*X)+Plateau). B) Extracted values from Pmm2-GFP depletion kinetics fitted against one-phase exponential decay model. hpf, hours post fertilization; NAA, auxin.
